# Coupling Flux Balance Analysis with Reactive Transport Modeling through Machine Learning for Rapid and Stable Simulation of Microbial Metabolic Switching

**DOI:** 10.1101/2023.02.06.527371

**Authors:** Hyun-Seob Song, Firnaaz Ahamed, Joon-Yong Lee, Christopher C. Henry, Janaka N. Edirisinghe, William C. Nelson, Xingyuan Chen, J. David Moulton, Timothy D. Scheibe

## Abstract

Integration of genome-scale metabolic networks with reactive transport models (RTMs) is an advanced simulation technique that enables predicting the changes of microbial growth and metabolism in space and time. Despite promising demonstrations in the past, computational inefficiency has been pointed out as a critical issue to overcome because it requires repeated implementation of linear programming (LP) to get flux balance analysis (FBA) solutions in every time step and every spatial grid. To address this challenge, we propose a new simulation method where we train/validate artificial neural networks (ANNs) using FBA solutions randomly sampled and incorporate the resulting reduced-order FBA model (represented as algebraic equations) into RTMs as source/sink terms. We demonstrate the efficiency of our method via a case study of *Shewanella oneidensis* MR-1 strain. During the aerobic growth on lactate, *S. oneidensis* produces metabolic byproducts (such as pyruvate and acetate), which are subsequently consumed as alternative carbon sources when the preferred ones are depleted. Simulating such intricate dynamics posed a considerable challenge, which we overcame by adopting the cybernetic approach that describes metabolic switches as the outcome of dynamic competition among multiple growth options. In both zero-dimensional batch and one-dimensional column configurations, the ANN-based reduced-order models achieved substantial reduction of computational time by several orders of magnitude compared to the original LP-based FBA models. Importantly, the ANN models produced robust solutions without any special measures to prevent numerical instability. These developments significantly promote our ability to utilize genome-scale networks in complex, multi-physics, and multi-dimensional ecosystem modeling.

## Introduction

Flux balance analysis (FBA) using genome-scale metabolic networks has proven to be a valuable tool in modeling microbial metabolism by providing a holistic view of an organism’s metabolic capabilities (Orth et al., 2010; Antoniewicz, 2015). In environmental sciences, these models have been applied in studying the interactions between microorganisms and their environment (Henry et al., 2016; Roy Chowdhury et al., 2019; Saifuddin et al., 2019; McClure et al., 2022). The implementation of genome-scale metabolic networks in reactive transport models (RTMs) is of particular interest, because it allows us to simulate dynamic changes in microbial metabolism together with the movement of fluids and solutes in space and time (Harcombe et al., 2014; Song et al., 2014; Phalak et al., 2016, 2022; Bauer et al., 2017; Borer et al., 2019). With the advent of high-throughput omics data, there is an increasing interest in using metabolic network models in ecosystem modeling as current simplistic biogeochemical models lack a high resolution required for effective integration of molecular data (Song et al., 2020).

Coupling FBA with RTMs poses significant computational challenges because it requires iterative implementation of linear programming (LP) in every time step and spatial grid. To avoid this issue, Scheibe and colleagues proposed indirect and direct coupling methods (Scheibe et al., 2009; Fang et al., 2011). Indirect coupling developed by Scheibe et al. (2009) generates a look-up table in advance by collecting a large set of FBA solutions in various possible environmental conditions, which is then referenced throughout the dynamic simulation of the RTM. However, not all FBA solutions pre-generated as such are relevant for reactive-transport simulations, indicating that the look-up table is likely to unnecessarily include a significant portion of FBA solutions that are never used by the RTM. In the direct coupling method proposed by Fang et al. (2011), a look-up table is quite small in the beginning or does not exist but progressively grows during dynamic simulations by adding new FBA solutions to the table if unavailable yet. Despite promising demonstrations in case studies, these approaches including indirect and direct coupling are not readily extended to large-scale, spatially heterogeneous systems with multiple chemical and biological species. Beside the painful process of building huge-size look-up tables, iterative communications between reactive transport simulators and the memory space (storing the FBA look-up table) impede efficient simulations, requiring more efficient approaches.

In this work, we propose a new method that enables coupling FBA with RTM in a computationally efficient way using machine learning. Instead of building a look-up table, we train artificial neural networks (ANNs) from randomly sampled FBA solutions, which serves as a reduced-order FBA model. As the resulting ANNs are represented as algebraic equations, they can be directly incorporated into source/sink terms in the RTM. We apply this idea to model dynamic growth of *Shewanella oneidensis* MR-1 strain (Pinchuk et al., 2010; Song et al., 2013), which exhibits complex growth patterns caused by dynamic metabolic switches among multiple substrates: in aerobic growth on lactate, the organism produces acetate and pyruvate as metabolic products; when lactate is depleted, it switches the carbon source over to pyruvate and produces acetate; when pyruvate is depleted, it takes up acetate for further growth. No computational methods have been formulated as of yet that would allow for the simulation of such intricate dynamics by joining FBA (or its surrogate models) and RTM. Because metabolic products in the previous stage become carbon sources for growth in the next stages, it is uncertain how to constrain the metabolic network model in dynamically varying media conditions. Training accurate ANNs that are able to portray such diverse metabolic traits of microorganisms is therefore an additional challenge. We demonstrate how these issues can be handled by using new computational methods proposed in this work. The performance of our methods is evaluated through case studies of the growth of *S. oneidensis* MR-1 in zero-dimensional batch and one-dimensional column reactors.

## Methods and Materials

### Metabolic network model of *S. oneidensis* MR-1

We used a genome-scale metabolic network of *S. oneidensis* MR-1 strain (termed iMR799) in the literature (Ong et al., 2014). The version of the model we used was loaded into KBase, which involved translating all model reactions and compounds to ModelSEED IDs. This was done to ensure that the iMR799 model would be interoperable with other KBase models as the ultimate goal of this effort is to simulate multi-species communities. The model in KBase was modified in several ways to ensure proper function: (1) three reactions ‘R_EX_TWEEN_20_E’, ‘R_EX_GEL_E’, and ‘R_EX_CAS_E’ were removed from the model as these allowed large mass influx without balance within KBase; (2) the “glycogen” metabolite in the model was adjusted to “glycogenmonomer” to ensure it does not collide with multimer versions of glycogen in the KBase namespace (such a collision results in mass imbalanced reactions in the translated model); (3) we revised the biomass production equation by adding ATP hydrolysis that is missing in the original model, i.e., *c* ATP + *c* H_2_O ↔ *c* ADP + *c* Pi + *c* H^+^ where *c* is the stoichiometric coefficient of the compounds involved in this reaction. When the value of *c* is low, the model produces excessive ATP and there is no difference between aerobic vs. anaerobic growth. As a default value, we put 40.11 for *c* in the metabolic network model (i.e., the SBML file), which was later adjusted (along with other additional parameters) in FBA simulations such that the model fits the experimentally measured yield data. The resulting metabolic network is confirmed to produce biomass from three individual carbon sources (i.e., lactate, pyruvate, and acetate).

### Formulation of multi-step FBA

FBA is an LP-based method that enables determining a flux distribution in a metabolic network by maximizing biomass production (while any biologically appropriate metabolic objectives can also be considered). As a common option, FBA subsequently minimizes a sum of flux magnitudes as a secondary objective to reduce the variations of LP solutions. This typical implementation, however, failed to predict experimentally observed byproduct formation in *S. oneidensis*, i.e., the production of pyruvate and acetate during the growth on lactate; the production of acetate during the growth on pyruvate. To align FBA simulations with experimental observations, we designed a multi-step FBA that involves following a sequence of LPs.

For the simulation of growth on lactate (*Lac*), for example, we carried out the following three steps. First, we perform a typical FBA to determine the maximum rate of biomass (*Bio*) production, which is denoted by 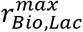 where the subscript after the comma (*Lac*) indicates what the carbon source is in the growth media. Second, we perform another round of LP to determine the maximum rate of pyruvate (*Pyr*) production (denoted by 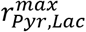) under an additional constraint: 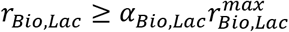 where *α*_*Bio*,*Lac*_ is a tuning parameter (ranging from 0 to 1) introduced to constrain the biomass production because otherwise the model does not produce enough pyruvate observed in the experiment. Lastly, we perform LP again to determine the maximum rate of acetate (*Ace*) production under the two additional constraints: 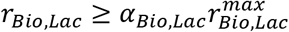 and 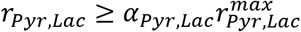 where *α*_*Bio*,*Lac*_ is the same parameter introduced in the previous step and *α*_*Pyr*,*Lac*_ is a new tuning parameter (ranging from 0 to 1) introduced to constrain the production of pyruvate.

In a similar fashion, we designed a two-step LP problem to simulate the growth with pyruvate as the sole carbon source. It should be clear that FBA in this case introduces a new parameter, *α*_*Bio*,*Pyr*_, which constrains the production of biomass. There is no need for a multi-step LP for simulating growth on acetate as no production of metabolic byproducts was experimentally detected. All FBA simulations were performed in minimal growth media conditions.

### Nonlinear optimization to determine the parameters in the multi-step FBA

The multi-step FBA formulated above includes a set of parameters to tune: *c* (i.e., the stoichiometric coefficient in the metabolic network model), *γ*_*Bio*,*Lac*_, *γ*_*Pyr*,*Lac*_, and *γ*_*Bio*,*Pyr*_. These four parameters were determined such that the error between predicted vs. measured yields of biomass and byproducts, i.e.,

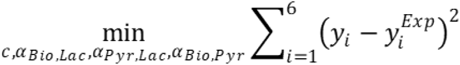

where *y_i_* ′*s* and 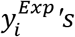 are FBA-predicted and experimentally-measured yields of biomass and metabolic products, respectively, where the subscripts 1 to 3 denote the yields of biomass, pyruvate, and acetate from the consumption of lactate, the subscripts 4 and 5 are the yields of biomass and acetate from the consumption of pyruvate, and the subscript 6 is the yield of biomass from the consumption of acetate.

We obtained experimentally measured yield data 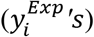 from Song et al. (2013). We calculated predicted yields (*y_i_* ′*s*) from multi-step FBA solutions (described in the previous section) by normalizing the production fluxes of biomass and byproducts with respect to the uptake flux of a given carbon source, i.e.,

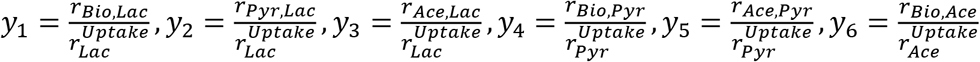

We solved this nonlinear optimization problem using Matlab solver ‘fminsearch.m’ to determine optimal values of the four parameters (*c*, *α*_*Bio*,*Lac*_, *α*_*Pyr*,*Lac*_, and *α*_*Bio*,*Pyr*_.). In doing this, we set the maximum magnitudes of the carbon uptake rate based on the kinetic parameters previously determined in Song et al. (2013), which are 22.1, 8.19, and 4.39 for lactate, pyruvate, and acetate, respectively.

### ANN model building

We built three ANN models as surrogate models of FBA to simulate the aerobic growth of *S. oneidensis* from lactate, pyruvate, and acetate, respectively. For each case, we generated 5,000 FBA solutions with different values of the uptake rates of oxygen and a given carbon source, which were randomly chosen by assuming uniform distributions over the range from zero to their max values determined by the reaction rate constants identified in a previous work (Song et al., 2013). We split the dataset (i.e., 5,000 FBA solutions) into training (70%), validation (15%) and testing (15%) subsets. Each ANN takes the upper limits of the magnitudes of uptake rates of oxygen and a given carbon source as inputs and predicts the production rates of biomass and metabolic byproducts. We also included the ‘actual’ uptake rate of oxygen as part of output variables because the imposed upper limit and actual uptake for oxygen were not necessarily the same (see **Fig. S1**). We determined the optimal numbers of nodes and layers in each ANN as key hyperparameters. Through the grid search, we determined the minimum numbers of nodes and layers, beyond which no measurable improvement in the accuracy of output variables was observed. We used the Bayesian regularization backpropagation method for optimization. We performed all calculations required for developing ANN models using the neural network toolbox in the Matlab 2022b version.

## Results

### Characterization of the FBA solution space

Our multi-step LP ensured the FBA model of *S. oneidensis* to produce metabolic byproducts as experimentally observed. As described in **Methods and Materials**, the multi-step LP formulation contains a set of parameters: *c*, *α*_*Bio*,*Lac*_, *α*_*Pyr*,*Lac*_, and *α*_*Bio*,*Pyr*_ that need to be optimized, where the constant *c* is the stoichiometric coefficient of ATP in the biomass production equation, which is a metric for the growth-associated maintenance, while the other three denote the fractional production of metabolic byproducts compared to their theoretical maximum values under given constraints. Accurate determination of these values is critical for robust dynamic simulation of complex growth patterns of *S. oneidensis*. By performing nonlinear optimization, we determined *c* to be 195.45, which was consistent with the value estimated in a previous study in the literature by Pinchuk et al. (2010) (= 220.22), and the other parameters were identified as follows: *α*_*Bio*,*Lac*_ = *0*.*6721*, *α*_*Pyr*,*Lac*_ = *0*.*6848*, *α*_*Bio*,*Pyr*_ = *0*.*6837*, indicating the actual production of all metabolic byproducts in the model to be below 70% of their full capacity in all cases.

With these parameter values, we performed the multi-step LP algorithm to characterize FBA-predicted exchange fluxes that are relevant for simulating metabolic switches in *S. oneidensis* and are later used as input data for building ANNs. The exchange fluxes of interest include the uptake rates of oxygen and individual carbon sources (lactate, pyruvate, or acetate), and the production rates of biomass and metabolic byproducts (i.e., the production of pyruvate and acetate from the consumption of lactate, or the production of acetate from the consumption of pyruvate). For each of the carbon sources that are utilizable by *S. oneidensis* for growth, we individually visualized FBA-estimated exchange fluxes of substrate consumption (i.e., carbon and oxygen uptake) (**Fig. S1**) and biomass production (**Fig. 1**) and byproduct production (**Fig. 2**). The fluxes of biomass production were characterized by the three phase planes (differentiated with dashed lines), each of which represented carbon-, oxygen- and both C&O-limited growth conditions, respectively (**Fig. 1**). The actual uptake rates realized by FBA were the same as the imposed upper limits for carbon sources, which was however not the case for oxygen (**Fig. S1**), indicating that, in addition to the production rates of biomass and metabolic products, it is necessary to store the values of oxygen uptakes to inform ANNs for predictions.

**Figure 1:**
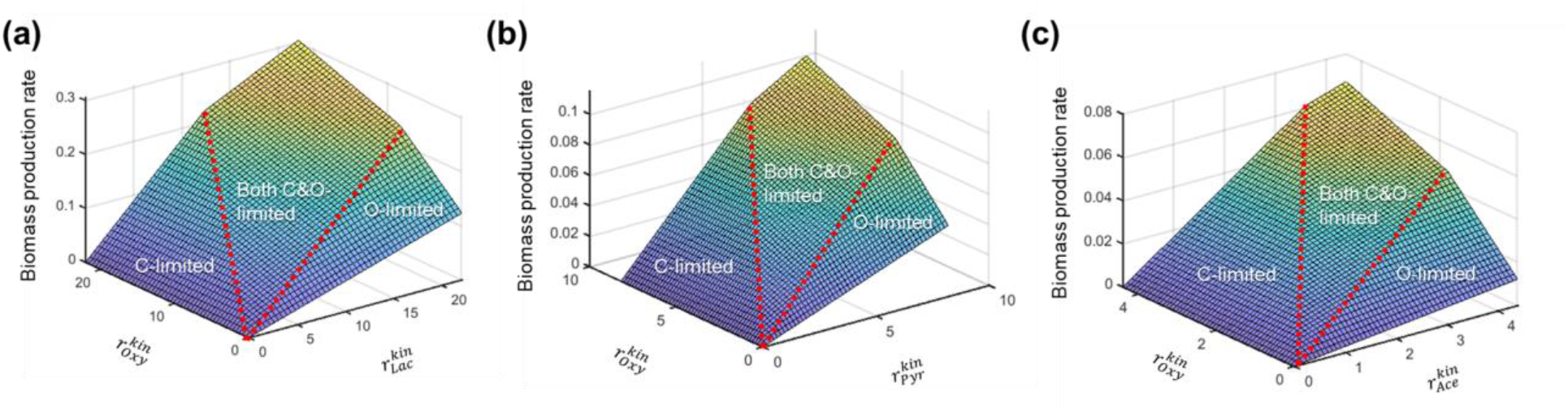
FBA prediction of biomass production rates in *S. oneidensis* MR-1: the aerobic growth on (a) lactate, (b) pyruvate, and (c) acetate. Three phase planes differentiated by dashed lines in each panel denote the growth limited by carbon (C), oxygen (O), and both C&O, respectively.

**Figure 2:**
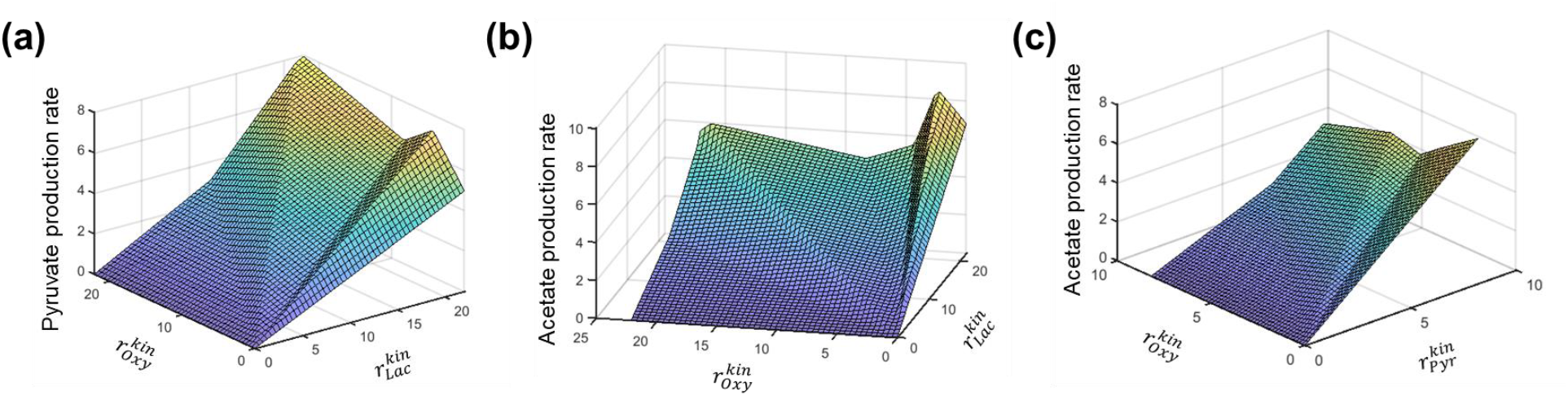
FBA prediction of by-product production rates in *S. oneidensis* MR-1: (a) pyruvate production rate from lactate consumption, (b) acetate production rate from lactate consumption, and (b) acetate production rate from pyruvate consumption.

Patterns for the production rates of metabolic byproducts in **Fig. 2** were much more complex than the production rate of biomass in **Fig. 1**. The numbers of phase planes for the former were six and seven for the production of pyruvate and acetate from lactate, and four for the production of acetate from pyruvate. Highly nonlinear behaviors observed in the response curves in **Fig. 2** may be ascribed to the implementation of multi-step LP with a set of parameters introduced to constrain the production rates of metabolic byproducts, posing a challenge in building quantitative ANN models in predicting their fluxes.

### Robust performance of ANNs: comparison of single- vs. multi-output models

We developed ANNs as surrogate models of FBA to predict the growth of *S. oneidensis* on lactate, pyruvate, and acetate, respectively. For each of these carbon sources, we compared the following two approaches: (1) we generated a set of ANNs, each of which aims to predict only a single output flux by taking two input fluxes (i.e., upper limits of the carbon substrate and oxygen uptakes) and is therefore a multi-input single-output (MISO) model; (2) we also built a single ANN to predict all exchange fluxes, which is a multi-input multi-output (MIMO) model. Similarly, we built MISO and MIMO models for the growth from pyruvate and acetate, respectively. We performed the grid search to determine the optimal number of nodes and layers.

In the case with lactate as the carbon source, the numbers of nodes and layers in the MISO models varied from six to ten (nodes), and two to three (layers) (**Table S1**). All MISO models with lactate as the carbon source led to the high correlations (> 0.9999) between the target values (i.e. FBA solutions) and ANN outputs in testing, validation, and testing. Interestingly, the MIMO model achieved the equivalent performance with slightly larger numbers of nodes and layers (i.e., 10 and 5, respectively) compared to the MISO models. This observation was consistent in other cases with pyruvate and acetate as the carbon sources. For convenience, therefore, we used the MIMO models throughout all simulations in this paper.

### Simulation of the metabolic switching of *S. oneidensis* in a batch culture

We demonstrate the effectiveness of ANNs as surrogate FBA models via the case study of the aerobic growth of *S. oneidensis* in a batch reactor in this section and in a one-dimensional column reactor in the next section. Mass balance equations for the batch growth are given a set of ordinary differential equations (**Table 1**). Dynamic FBA (dFBA) is the method developed to integrate FBA with microbial growth models (i.e., ODEs) (Mahadevan et al., 2002). While typical implementation of dFBA is conceptually simple, simulating metabolic switches among alternative carbon sources as in our case poses a challenge because the information on substrate concentrations and kinetics alone cannot tell whether the corresponding compounds will be consumed or produced because this is the outcome of metabolic regulation in microorganisms.

**Table 1.**
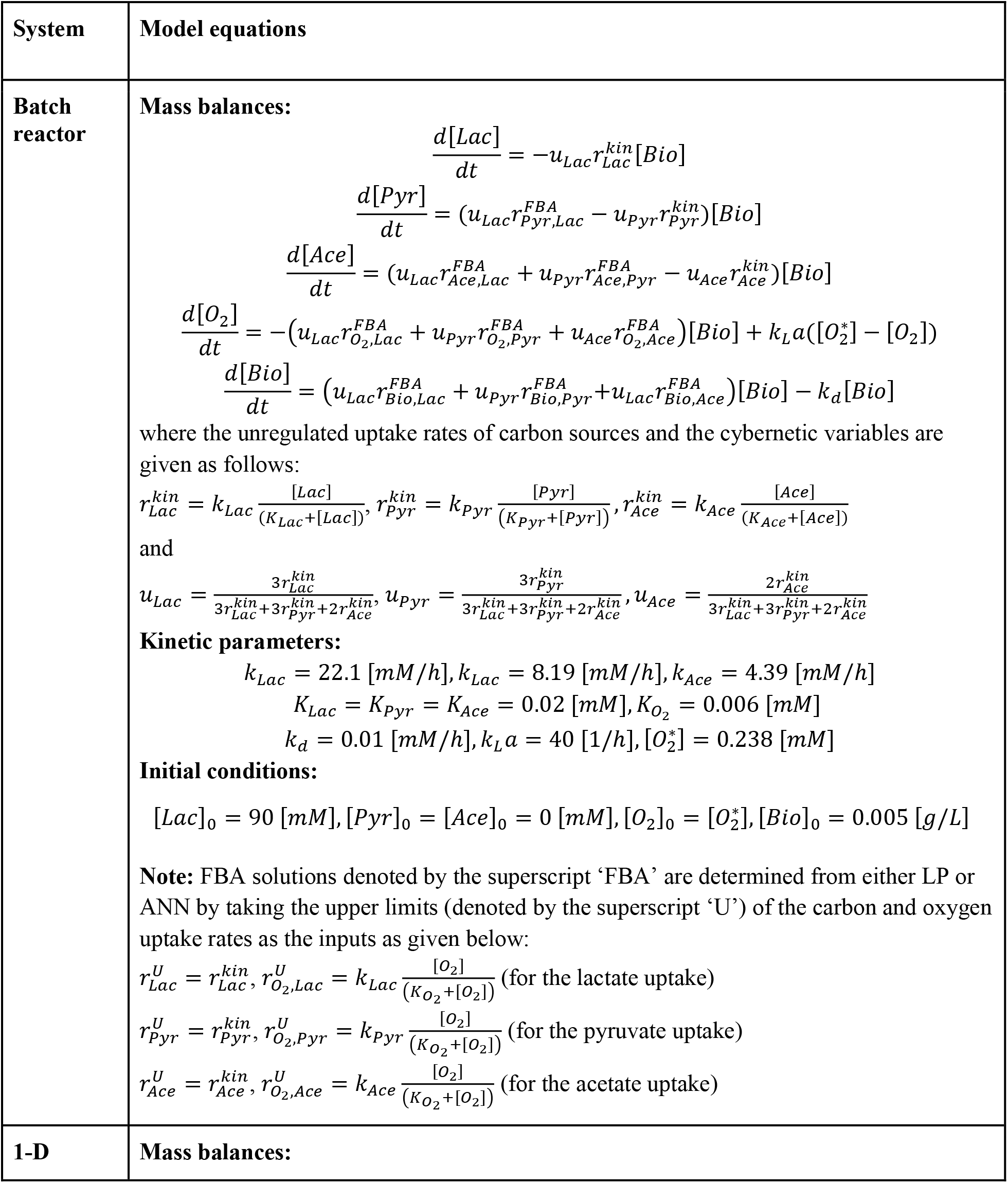
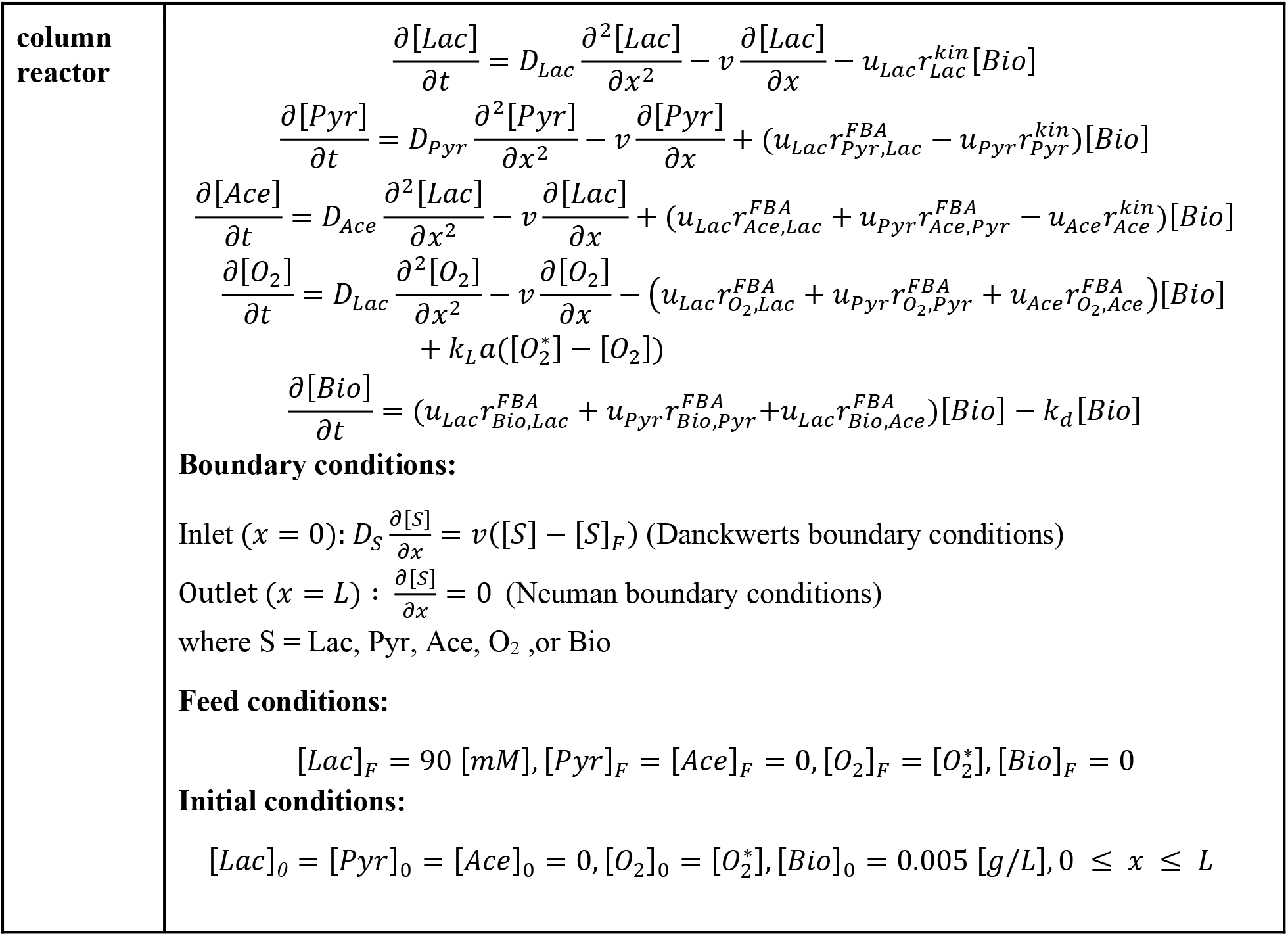
Mass balance equations for simulating the aerobic growth of *S. oneidensis* MR-1 on lactate in batch and one-dimensional column reactors. The symbol [ ] is used to denote concentration of substrates and biomass, *r*’s are fluxes and the subscript ‘FBA’ denotes fluxes estimated by FBA, *k_L_a* is the volumetric coefficient of oxygen transfer between the growth media and atmosphere, and [*O*\*_*2*_] is the saturated level of oxygen at the standard condition.

While critical for robust simulation of metabolic switching, the incorporation of metabolic regulation is not straightforward because its full mechanistic details are generally not known. We therefore used the cybernetic approach as a key component of our model, which provides a rational mathematical description of dynamic regulation based on an optimal control theory without introducing additional parameters (Ramkrishna and Song, 2012, 2018). By viewing the metabolism of *S. oneidensis* as a dynamic combination of three FBA solutions obtained from each of the carbon sources, we independently performed FBA three times in each time step to get the flux vector in *S. oneidensis* associated with the consumption of lactate, pyruvate, and acetate, respectively. The resulting three flux vectors were subsequently combined in proportion to their contribution to promote a pre-defined metabolic objective, for which we used ‘carbon’ uptake rate in this work. This combination is realized through the cybernetic variables *u_Lac_*, *u_Pyr_*, and *u_Ace_* as defined in **Table 1** where the numerical values multiplied on the uptake rates denote the number of carbon contained in each carbon source. FBA solutions are quickly generated using ANNs developed in the previous section.

In **Fig. 3**, we presented the resulting simulations of aerobic growth of *S. oneidensis* in the batch reactor by comparing the original LP- and ANN-based dFBA models. While both models showed consistency in simulating dynamic metabolic switches through the sequential utilization of alternative carbon sources, the computational time taken by the ANN-based dFBA was only 0.1 to 0.2% compared to that of the original dFBA. Notably, the ANN-based dFBA model showed numerical stability throughout dynamic simulations, while the original dFBA was stuck when the level of any of the carbon sources was near to zeros. While this issue occurring for the original dFBA may be relieved by using specially designed numerical solvers (Chen et al., 2016; Phalak et al., 2016), no special treatment was necessary for the ANN-based dFBA model.

**Figure 3:**
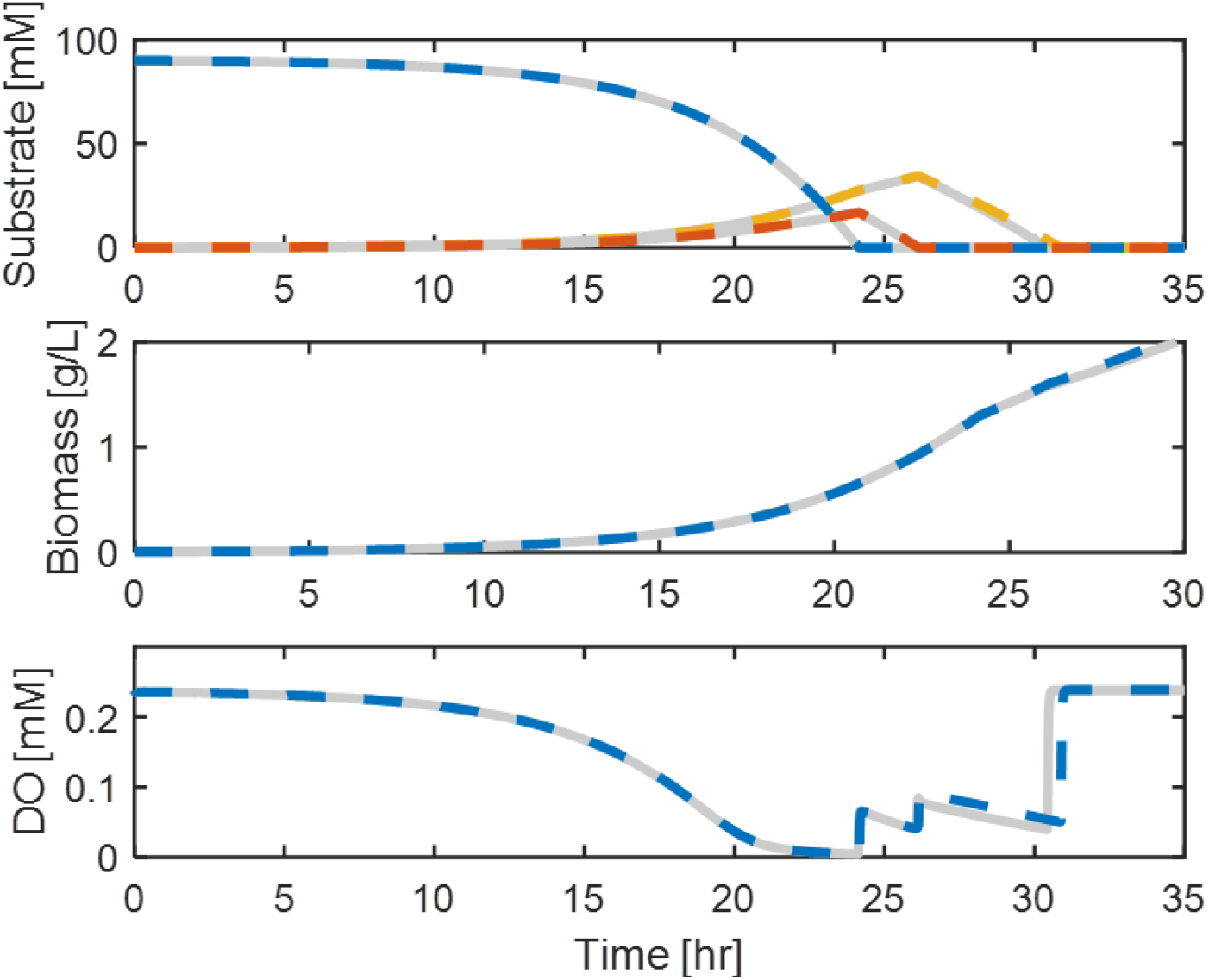
Comparison of FBA simulations of aerobic growth of *S. oneidensis* MR-1 in a batch reactor using LP (solid lines) and ANN models (dashed lines).

### Simulation of *S. oneidensis* growth in a one-dimensional column reactor model though FBA-RTM coupling

We extend the method to couple FBA (or ANN) with microbial growth models to RTMs where the concentrations of substrates and biomass are spatially distributed. For simplicity, we consider a one-dimensional column reactor in this work. The mass balances of this column model are given as a set of partial differential equations (PDEs) (**Table 1**). To solve PDEs, we discretized the spatial derivative terms into algebraic forms, while keeping the time derivative terms as is. This conversion termed ‘the method of lines’ allows us to use any ODE solvers to simulate PDE models. As a more notable advantage of using this conversion method, we can also use the same simulation technique employed in the previous section to simulate the metabolic switching of *S. oneidensis* in the batch reactor. No additional special treatment is required for coupling FBA and RTM.

For the one-dimensional column reactor with 100 spatial grids, our ANN-based column model generated the solutions within one minute on a personal desktop computer. We could not make a direct comparison with the original LP-RTM model because the latter failed to generate stable numerical solutions with the same number of grids. Instead, we were able to obtain the solutions from the original model only for the configurations with low grid densities (i.e., only up to the 30 number of grids). Through the comparison using these coarse-grained simulations with the grid number of 10, 20, and 30, we found that the reduction of computational time by the ANN-based RTM was about three orders of magnitude (**Fig. 4**), a similar performance we observed in the previous batch simulations. For the models with higher grid densities, therefore, we provided only estimated computational time for the original LP-RTM. Again, this result highlights that numerical stability is an additional key advantage using the ANN-RTM.

**Figure 4:**
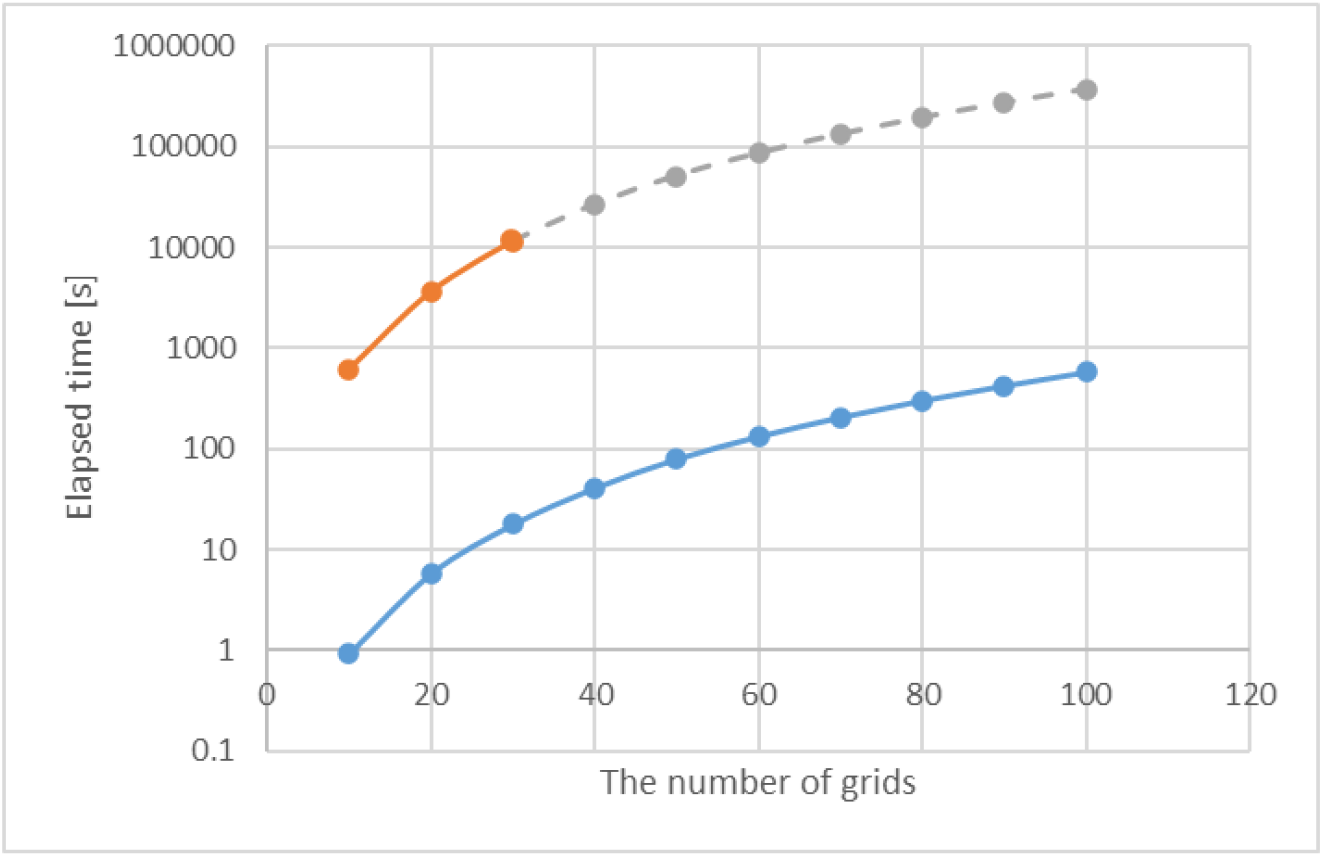
Comparison of simulation times in a one-dimensional column reactor as a function of axial grid size: FBA simulation times by the ANN model (blue solid line) and LP (orange solid line). Gray dashed line denotes ‘expected’ times by LP (for 40 to 100 grids), which is estimated by multiplying an average ratio of the two results for the first three results (for 10 to 30 grids).

## Discussion

Our new computational method significantly expands the scope of genome-scale metabolic network modeling to predict microbial growth in space and time by coupling with RTMs. By using machine learning models (i.e., ANNs) as the source/sink terms in RTMs instead of solving LP iteratively, our method drastically reduced computational time, while providing robust and stable simulations by eliminating numerical errors commonly encountered when solving FBA-coupled ordinary or partial differential equations. Due to these promising properties, the ANN-based surrogate FBA model developed in this work is currently incorporated as a key component of *CompLab*, a recently developed Lattice Boltzmann (LB)-based modeling tool to simulate fluid flow and solute transport in porous media by coupling metabolic networks with RTMs (Jung et al., 2022).

Our development also opens up new opportunities for modeling the intricate metabolic switching among multiple alternative carbon sources, a common phenomenon observed in metabolic systems. While previous studies in the literature have already demonstrated the capability of simulating dynamic metabolic changes in varying environments by coupling FBA with RTMs, our method is the first to enable FBA for the simulation of drastic shifts in metabolism, e.g., as those observed in *S. oneidensis*. This type of simulation is not readily implementable by typical kinetic description of uptake rates because in our case the same carbon source can be consumed as the carbon source for growth or produced as a byproduct depending on the context. Inspired by the work by Song et al. (2013), we resolved this issue by using the cybernetic modeling approach where we describe the dynamic shifts in *S. oneidensis* as the outcome of competition among three FBA solutions associated with three individual carbon sources, respectively. The cybernetic modeling was originally developed as an independent tool for simulating complex growth patterns of microorganisms (Ramkrishna and Song, 2012, 2018) and microbial communities (Song and Liu, 2015; Song et al., 2017, 2018, 2020), the effectiveness of which was greatly enhanced when used in conjunction with metabolic pathways and networks using elementary flux mode analysis (Kim et al., 2008, 2012; Young et al., 2008; Song et al., 2009; Song and Ramkrishna, 2010, 2011, 2012; Franz et al., 2011) as well as FBA (Vilkhovoy et al., 2016; Luo et al., 2023). The present work serves as an outstanding example in this direction.

While critically important in making FBA predictions consistent with experimental observations in our work, the introduction of tuning parameters (such as the constraints on the production of biomass and metabolic byproducts in metabolism of *S. oneidensis*) may be considered an *ad hoc* remedy because those parameters are determined through datafit without having a sufficient biological basis. Imposing additional constraints on the internal pathways in a more mechanistic way may enable the metabolic network model of *S. oneidensis* to produce metabolic byproducts as experimentally observed without manual forcing. Further development of metabolic networks of *S. oneidensis* in this direction is an important task in the future, while a major focus in this work was to develop and demonstrate the power of machine learning as a reduced-order FBA model in simulating RTMs.

Overall, the use of machine learning to achieve the reduction of computational time in metabolic network modeling by orders of magnitude represents a significant advance in diverse disciplines, particularly in the field of systems biology, microbial ecology, biogeochemistry and even earth science, where FBA is being used and can serve as a prime modeling tool. We envision that this capability will become more important in the future as the application of FBA is being extended to increasingly more complex systems. In systems biology, for example, FBA is used to model the interactions among multiple human organs by integrating the metabolic networks of each organ into a larger, whole-body model (Bordbar et al., 2011; Thiele et al., 2020). Similarly, the utility of FBA could be extended to predict interspecies metabolic interactions and cross-feeding in complex environmental microbial communities by integrating individual metabolic networks. The greatest potential benefit of this method is expected to be seen in ecosystem modeling, as it allows for the integration of genome-scale metabolic networks with RTMs, which govern the movement of chemical and biological species. In order to further improve this method, an important next step is to develop machine learning techniques to effectively incorporate multi-omics data (such as genomics, transcriptomics, proteomics, and metabolomics) into genome-scale metabolic networks and to couple them with RTMs. This will enhance the ability to understand and link molecular-level mechanisms to large-scale system dynamics, which will accelerate new scientific discoveries.

## Funding

This research was supported by the U.S. Department of Energy, Office of Science, Office of Biological and Environmental Research, Environmental System Science (ESS) Program. This contribution originates from the River Corridor Scientific Focus Area (SFA) project at Pacific Northwest National Laboratory (PNNL) and was supported by the partnership with the IDEAS-Watersheds. Pacific Northwest National Laboratory is operated by Battelle Memorial Institute for the U.S. Department of Energy under Contract DE-AC05-76RL01830.

## Conflict of Interest

The authors declare that the research was conducted in the absence of any commercial or financial relationships that could be construed as a potential conflict of interest.

---

**Supplementary Table S1.** The number of nodes and layers in ANN models. The two numbers in the parenthesis respectively denote the numbers of nodes and layers chosen for predicting output variables. MISO = multi-input single-output, and MIMO = multi-input multi-output.

| Input variables | Output variables in MISO models |  |  |  | Output variables in MIMO models |
| --- | --- | --- | --- | --- | --- |
|  | Oxygen uptake rate | Biomass production rate | Pyruvate production rate | Acetate production rate | All |
| Upper limit to the uptake rates of lactate and oxygen | (6,2) | (6,2) | (8,2) | (10,3) | (10,5) |
| Upper limit to the uptake rates of pyruvate and oxygen | (6,2) | (4,2) | N/A | (6,2) | (8,4) |
| Upper limit to the uptake rates of acetate and oxygen | (4,2) | (6,2) | N/A | N/A | (6,2) |

**Supplementary Figure S1:**
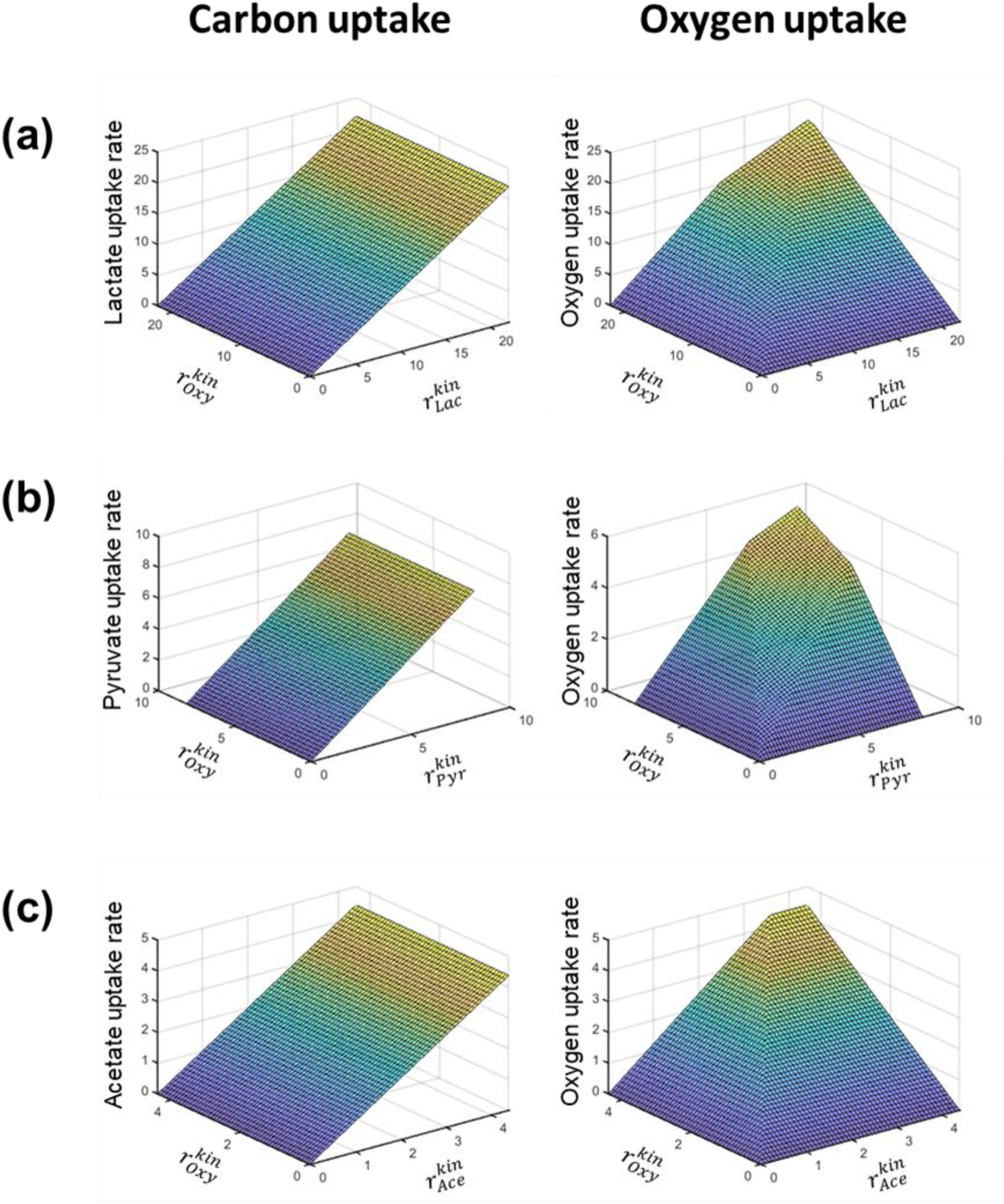
Carbon source-specific FBA-determined carbon uptake rates (left panels) and oxygen uptake rates (right panels) in the growth of *S. oneidensis* MR-1. Carbon source: (a) lactate, (b) pyruvate, and (c) acetate.

## Notes

### Competing Interest Statement

The authors have declared no competing interest.

## References

Antoniewicz, M. R. (2015). Methods and advances in metabolic flux analysis: a mini-review. J. Ind. Microbiol. Biotechnol. 42, 317–325. doi: 10.1007/s10295-015-1585-x.

Bauer, E., Zimmermann, J., Baldini, F., Thiele, I., and Kaleta, C. (2017). BacArena: Individual-based metabolic modeling of heterogeneous microbes in complex communities. PLOS Comput. Biol. 13, e1005544. doi: 10.1371/journal.pcbi.1005544.

Bordbar, A., Feist, A. M., Usaite-Black, R., Woodcock, J., Palsson, B. O., and Famili, I. (2011). A multi-tissue type genome-scale metabolic network for analysis of whole-body systems physiology. BMC Syst. Biol. 5, 180. doi: 10.1186/1752-0509-5-180.

Borer, B., Ataman, M., Hatzimanikatis, V., and Or, D. (2019). Modeling metabolic networks of individual bacterial agents in heterogeneous and dynamic soil habitats (IndiMeSH). PLOS Comput. Biol. 15, e1007127. doi: 10.1371/journal.pcbi.1007127.

Chen, J., Gomez, J. A., Höffner, K., Phalak, P., Barton, P. I., and Henson, M. A. (2016). Spatiotemporal modeling of microbial metabolism. BMC Syst. Biol. 10, 21. doi: 10.1186/s12918-016-0259-2.

Fang, Y., Scheibe, T. D., Mahadevan, R., Garg, S., Long, P. E., and Lovley, D. R. (2011). Direct coupling of a genome-scale microbial in silico model and a groundwater reactive transport model. J. Contam. Hydrol. 122, 96–103. doi: 10.1016/j.jconhyd.2010.11.007.

Franz, A., Song, H.-S., Ramkrishna, D., and Kienle, A. (2011). Experimental and theoretical analysis of poly(β-hydroxybutyrate) formation and consumption in Ralstonia eutropha. Biochem. Eng. J. 55, 49–58. doi: 10.1016/j.bej.2011.03.006.

Harcombe, W. R., Riehl, W. J., Dukovski, I., Granger, B. R., Betts, A., Lang, A. H., et al. (2014). Metabolic resource allocation in individual microbes determines ecosystem interactions and spatial dynamics. Cell Rep. 7, 1104–1115. doi: 10.1016/j.celrep.2014.03.070.

Henry, C. S., Bernstein, H. C., Weisenhorn, P., Taylor, R. C., Lee, J.-Y., Zucker, J., et al. (2016). Microbial Community Metabolic Modeling: A Community Data-Driven Network Reconstruction. J. Cell. Physiol. 231, 2339–2345. doi: 10.1002/jcp.25428.

Jung, H., Song, H.-S., and Meile, C. (2022). CompLaB v1.0: a scalable pore-scale model for flow, biogeochemistry, microbial metabolism, and biofilm dynamics. EGUsphere, 1–21. doi: 10.5194/egusphere-2022-1016.

Kim, J. I., Song, H.-S., Sunkara, S. R., Lali, A., and Ramkrishna, D. (2012). Exacting predictions by cybernetic model confirmed experimentally: Steady state multiplicity in the chemostat. Biotechnol. Prog. 28, 1160–1166. doi: 10.1002/btpr.1583.

Kim, J. I., Varner, J. D., and Ramkrishna, D. (2008). A hybrid model of anaerobic E. coli GJT001: Combination of elementary flux modes and cybernetic variables. Biotechnol. Prog. 24, 993–1006. doi: 10.1002/btpr.73.

Luo, H., Li, P., Ji, B., and Nielsen, J. (2023). Modeling the metabolic dynamics at the genome-scale by optimized yield analysis. Metab. Eng. 75, 119–130. doi: 10.1016/j.ymben.2022.12.001.

Mahadevan, R., Edwards, J. S., and Doyle, F. J. (2002). Dynamic Flux Balance Analysis of Diauxic Growth in Escherichia coli. Biophys. J. 83, 1331–1340. doi: 10.1016/S0006-3495(02)73903-9.

McClure, R., Farris, Y., Danczak, R., Nelson, W., Song, H.-S., Kessell, A., et al. (2022). Interaction Networks Are Driven by Community-Responsive Phenotypes in a Chitin-Degrading Consortium of Soil Microbes. mSystems 7, e00372–22. doi: 10.1128/msystems.00372-22.

Ong, W. K., Vu, T. T., Lovendahl, K. N., Llull, J. M., Serres, M. H., Romine, M. F., et al. (2014). Comparisons of Shewanella strains based on genome annotations, modeling, and experiments. BMC Syst. Biol. 8, 31. doi: 10.1186/1752-0509-8-31.

Orth, J. D., Thiele, I., and Palsson, B. Ø. (2010). What is flux balance analysis? Nat. Biotechnol. 28, 245–248. doi: 10.1038/nbt.1614.

Phalak, P., Bernstein, H. C., Lindemann, S. R., Renslow, R. S., Thomas, D. G., Henson, M. A., et al. (2022). Spatiotemporal Metabolic Network Models Reveal Complex Autotroph-Heterotroph Biofilm Interactions Governed by Photon Incidences. IFAC-Pap. 55, 112–118. doi: 10.1016/j.ifacol.2022.07.430.

Phalak, P., Chen, J., Carlson, R. P., and Henson, M. A. (2016). Metabolic modeling of a chronic wound biofilm consortium predicts spatial partitioning of bacterial species. BMC Syst. Biol. 10, 90. doi: 10.1186/s12918-016-0334-8.

Ramkrishna, D., and Song, H.-S. (2012). Dynamic models of metabolism: Review of the cybernetic approach. AIChE J. 58, 986–997. doi: 10.1002/aic.13734.

Ramkrishna, D., and Song, H.-S. (2018). Cybernetic Modeling for Bioreaction Engineering. Cambridge University Press.

Roy Chowdhury, T., Lee, J.-Y., Bottos, E. M., Brislawn, C. J., White, R. A., Bramer, L. M., et al. (2019). Metaphenomic Responses of a Native Prairie Soil Microbiome to Moisture Perturbations. mSystems 4, e00061–19. doi: 10.1128/mSystems.00061-19.

Saifuddin, M., Bhatnagar, J. M., Segrè, D., and Finzi, A. C. (2019). Microbial carbon use efficiency predicted from genome-scale metabolic models. Nat. Commun. 10, 3568. doi: 10.1038/s41467-019-11488-z.

Scheibe, T. D., Mahadevan, R., Fang, Y., Garg, S., Long, P. E., and Lovley, D. R. (2009). Coupling a genome-scale metabolic model with a reactive transport model to describe in situ uranium bioremediation. Microb. Biotechnol. 2, 274–286. doi: 10.1111/j.1751-7915.2009.00087.x.

Song, H.-S., Cannon, W. R., Beliaev, A. S., and Konopka, A. (2014). Mathematical modeling of microbial community dynamics: a methodological review. Processes 2, 711–752.

Song, H.-S., and Liu, C. (2015). Dynamic Metabolic Modeling of Denitrifying Bacterial Growth: The Cybernetic Approach. Ind. Eng. Chem. Res. 54, 10221–10227. doi: 10.1021/acs.iecr.5b01615.

Song, H.-S., Morgan, J. A., and Ramkrishna, D. (2009). Systematic development of hybrid cybernetic models: Application to recombinant yeast co-consuming glucose and xylose. Biotechnol. Bioeng. 103, 984–1002. doi: 10.1002/bit.22332.

Song, H.-S., and Ramkrishna, D. (2010). Prediction of metabolic function from limited data: Lumped hybrid cybernetic modeling (L-HCM). Biotechnol. Bioeng. 106, 271–284. doi: 10.1002/bit.22692.

Song, H.-S., and Ramkrishna, D. (2011). Cybernetic models based on lumped elementary modes accurately predict strain-specific metabolic function. Biotechnol. Bioeng. 108, 127–140. doi: 10.1002/bit.22922.

Song, H.-S., and Ramkrishna, D. (2012). Prediction of dynamic behavior of mutant strains from limited wild-type data. Metab. Eng. 14, 69–80. doi: 10.1016/j.ymben.2012.02.003.

Song, H.-S., Stegen, J. C., Graham, E. B., Lee, J.-Y., Garayburu-Caruso, V. A., Nelson, W. C., et al. (2020). Representing Organic Matter Thermodynamics in Biogeochemical Reactions via Substrate-Explicit Modeling. Front. Microbiol. 11, 531756. doi: 10.3389/fmicb.2020.531756.

Song, H.-S., Thomas, D. G., Stegen, J. C., Li, M., Liu, C., Song, X., et al. (2017). Regulation-Structured Dynamic Metabolic Model Provides a Potential Mechanism for Delayed Enzyme Response in Denitrification Process. Front. Microbiol. 8. Available at: https://www.frontiersin.org/articles/10.3389/fmicb.2017.01866 [Accessed February 4, 2023].

Song, X., Chen, X., Stegen, J., Hammond, G., Song, H.-S., Dai, H., et al. (2018). Drought Conditions Maximize the Impact of High-Frequency Flow Variations on Thermal Regimes and Biogeochemical Function in the Hyporheic Zone. Water Resour. Res. 54, 7361–7382. doi: 10.1029/2018WR022586.

Thiele, I., Sahoo, S., Heinken, A., Hertel, J., Heirendt, L., Aurich, M. K., et al. (2020). Personalized whole-body models integrate metabolism, physiology, and the gut microbiome. Mol. Syst. Biol. 16, e8982. doi: 10.15252/msb.20198982.

Vilkhovoy, M., Minot, M., and Varner, J. D. (2016). Effective Dynamic Models of Metabolic Networks. IEEE Life Sci. Lett. 2, 51–54. doi: 10.1109/LLS.2016.2644649.

Young, J. D., Henne, K. L., Morgan, J. A., Konopka, A. E., and Ramkrishna, D. (2008). Integrating cybernetic modeling with pathway analysis provides a dynamic, systems-level description of metabolic control. Biotechnol. Bioeng. 100, 542–559. doi: 10.1002/bit.21780.

